# Multi-Metric Quantitative MRI Identifies Spatially Distinct Age-Related Brain Differences in Female Bonnet Macaques

**DOI:** 10.1101/2025.04.09.648008

**Authors:** Laurel Dieckhaus, Kelsey McDermott, Jean-Philippe Galons, Carol Barnes, Elizabeth Hutchinson

**Affiliations:** University of Arizona

**Keywords:** Bonnet Macaque, Quantitative MRI, Diffusion MRI, Relaxometry, Tensor Based Morphometry, Support Vector Machine Classification

## Abstract

This study employs a data-driven, voxelwise analysis of high-resolution ex vivo quantitative MRI (qMRI) to examine age-related differences in brain morphometry and microstructure in female bonnet macaques. A binary classifier differentiated mid- and late-age groups, achieving the highest accuracy when integrating all MRI metrics rather than using diffusion or relaxometry alone. Diffusion-only and relaxometry-only classifiers revealed distinct, minimally overlapping spatial patterns, while the multi-metric approach captured a broader range of age-related differences. Tensor-based morphometry (TBM) differences were most pronounced in the neocortex, whereas the thalamus showed the highest classification accuracy despite minimal morphometric differences, suggesting unique tissue composition alterations. These findings highlight the complementary nature of diffusion, relaxometry, and morphometry qMRI metrics in aging research. Our results support the use of multi-parametric qMRI to identify age-vulnerable brain regions and highlight its potential for improving qMRI biomarkers in larger, longitudinal aging studies.

## 1. Introduction

Quantitative MRI (qMRI) metrics are sensitive to a gamut of brain tissue composition and microstructural features and to cellular and macromolecular alterations, making them potentially powerful biomarkers. However, the uniqueness and redundancy of biological information carried by different qMRI metrics across the brain has not been well established, as the majority of studies either compare metrics in targeted anatomical regions or evaluate a single MRI technique across the whole brain^1-3^. Although qMRI encompasses a wide range of techniques - from volumetrics and morphometry to microstructural MRI via diffusion to macromolecular and susceptibility mapping with relaxometry - the outcome metrics from different qMRI frameworks may be interdependent due to their shared MRI signal dependence or the influence of co-occurring biological changes. Consequently, the utility of qMRI metrics for brain characterization or disease diagnosis would be increased with a more complete and unbiased understanding of the relative value and complementarity across the spectrum of available and promising qMRI metrics. The capability to perform bias-free, whole brain, multivariate analysis has been improved with the development of machine learning tools including support vector machine (SVM) classification^4^. SVM techniques can simultaneously indicate the predictive accuracy of single and grouped metrics on a voxelwise basis and in this study, we have applied SVM to evaluate a range of qMRI metrics for their age-dependence in the healthy brain.

In the context of aging, qMRI metrics could serve as essential tools for understanding normal, age-dependent brain changes and for detecting abnormalities before cognitive symptoms emerge. Cross-sectional and longitudinal volumetric studies in mid- and late-aged adults have characterized the spatiotemporal patterns of brain volume loss during aging ^5-7^ and reduced hippocampal volume has even become a surrogate marker for neurodegenerative diseases^8^. However, volume and morphometry metrics are limited to reporting gross anatomic changes such as tissue gain or loss, and fail to capture subtle compositional changes that may precede atrophy^9-13^. Combining volumetrics with other quantitative MRI metrics is expected to provide a richer understanding of tissue environment changes on both macrostructural and microstructural scales^11^. In particular, diffusion and relaxometry qMRI can measure microstructural and compositional changes within each voxel^14-16^.

Among the different qMRI that leverage the relationship between MR signal and tissue environment, diffusion MRI has emerged as a prominent microstructural imaging method, especially in the context of both grey and white matter aging^14-19^ by its capability to probe length scale and shape features at the cellular scale. By comparison, relaxometry MRI provides metrics that are more influenced by the chemical composition of tissue within a voxel than by microstructural geometry effects ^11,20-22^. For example, the extremely short T2 relaxation component of water trapped between myelin sheaths (10-40 ms) can be estimated to report myelin content as Myelin Water Fraction (MWF),^22-29^ while the common DTI metric of fractional anisotropy (FA) measures the directional motion of water to probe the microstructural integrity of myelinated axons^11^. Each metric probes white matter state, but with different biologic underpinnings and applicability in aging studies. Within relaxometry techniques there is also a range of biological sensitivity from simple T2 mapping for intracellular and extracellular water content as well as the presence of iron^28,30,31^. More sophisticated techniques like Bound Pool Fraction (BPF) which uses magnetization transfer to probe macromolecular content ^22,32^ and susceptibility or R2* mapping of paramagnetic and diamagnetic materials like iron and calcium in brain tissue^14^.

A key challenge across all qMRI techniques, however, is a lack of specificity to underlying biology and interdependence of metrics that are influenced by the presence of multiple biochemical and microstructural factors. For instance, metrics like MWF and BPF, while considered markers for myelin, also retain signals from subcortical grey matter. Another example is that both iron content, macromolecular composition and structural boundaries (e.g., cell bodies and water content) can shorten T2 so that the underlying biology cannot be directly related to changes in quantitative T2 values. The low specificity commonly noted across most qMRI metrics underscores the need for comprehensive evaluation of multiple MRI metrics across tissue contexts to identify redundancy and specificity^10,33-35^. If this can be accomplished, the promise of combining diverse MRI metrics for a more complete view of tissue composition and microstructure will exceed the capabilities of single metric assessments.

Improving the understanding of metric correspondence to biological features and recognizing the strengths and limitations of different MRI approaches are crucial goals. While several quantitative MRI maps have demonstrated sensitivity as biomarkers for age-related brain changes, it remains unclear which maps are most sensitive to specific aging processes and whether different metrics provide complementary or redundant information ^10^. Traditional univariate approaches like General Linear Models (GLM) may fail to capture the combinatorial information offered by multi-parametric MRI^36^. Some multi-parametric approaches, such as Principal Component Analysis (PCA), transform features but complicate the extraction of original variables (e.g., specific MRI map types) ^37^. Linear Support Vector Machines (SVMs) are a supervised machine learning method that identifies a hyperplane which separates classes or categories using multivariate data^4^. SVMs have shown promise in contexts like cancer diagnosis ^38^ and recently in identifying white matter lesions in multiple sclerosis^39^. SVMs provide spatial identification of disease-related tissue by integrating multiple MRI maps, offering a holistic view of tissue composition^16^.

This study applied a binary SVM-based classifier to co-registered, multivariate qMRI metric maps of middle to old age (14-34y) bonnet macaque brains to identify predictive accuracy of qMRI metrics for age-related differences on a voxelwise basis. Using this data-driven approach, we aimed to **1) Identify regions most susceptible to aging at macrostructural and microstructural levels using qMRI** and **2) Highlight microstructural qMRI maps driving high-accuracy age classification predictions**. We selected the ex vivo bonnet macaque brain as a test bed for this study as non-human primates do not undergo the same age-related brain pathology as humans and provide an excellent model for studying lifespan-related brain differences^40^. Additionally, *ex vivo* MRI acquisition enabled high-resolution collection of many qMRI maps that would otherwise be time-constrained *in vivo*. We sought to identify age-related tissue changes by applying a voxel-wise SVM approach to identify both macrostructural and microstructural differences, while also characterizing the specific contributions of individual qMRI metrics to age-sensitive brain regions.

## 2. Methods

### 2.1 Samples and Sample Preparation

All specimens were obtained from a larger previous behavioral study^41^. Macaques were located and housed in Tucson, AZ facilities in temperature and humidity-controlled environments on a 12-hour light/dark cycle. Procedures were conducted in accordance with National Institutes of Health guidelines and the protocols were approved by the Institutional Animal Care and Use Committees at the University of Arizona. After transcardial perfusion, brains were refrigerated in 4% Paraformaldehyde and then transferred to a solution of PBS and sodium azide for longer than 2 weeks to ensure adequate rehydration before imaging. Brain samples received were of seven female bonnet macaques of ages 14-34 years (equivalent to 30– 75 human years; see Figure 1). These were categorized as middle age and late age based on the distribution of samples in the study. For MRI scanning, brains were placed in MR-compatible tube holders cushioned with medical gauze to minimize movement artifacts. The containers were filled with Flourinert (FC-3823, 3M, St. Paul, MN) and degassed using a vacuum chamber for approximately 2 hours prior to imaging.

**Figure 1:**
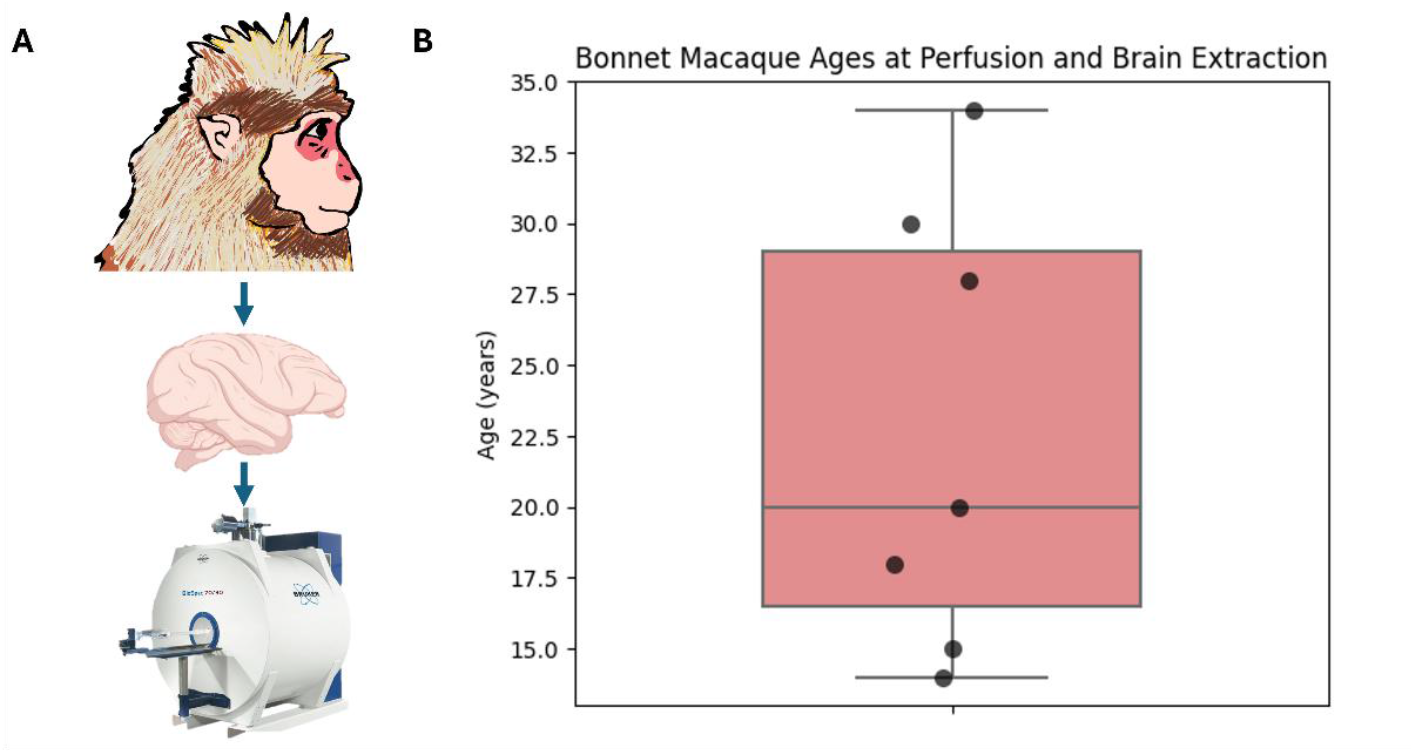
A) Cohort of female bonnet macaques were perfused and their brains were extracted and preserved in paraformaldehyde then transferred to a PBS and sodium azide solution. Extracted brains were individually imaged on a Pre-clinical 7T Bruker system. B) Age of female bonnet macaques at time of trancardial perfusion and brain extraction (age in years – y axis) with two separate distributions of ages defined as middle and late age

### 2.2 MRI Data Acquisition

Imaging was performed using a Bruker 7T scanner equipped with an 86mm quadrature coil and controlled by Paravision 360 software (versions 3.2 and 3.5). Five MRI scan types were collected using 3D acquisitions collected with coronal slices. For basic acquisition parameters, see Table 1. Sequence-specific parameters are as follows:

**Table 1:** MRI Acquisition Scan Types and Parameters.

| Maps Processed | High Resolution Anatomical | T2 and Myelin Water Fraction | T1 and Bound Pool Fraction (Quantitative Magnetization Transfer) | Diffusion Tensor | R2* |
| --- | --- | --- | --- | --- | --- |
| Scan Type | T1 FLASH | Multiple Spin Echo | Selective Inversion Recovery | Diffusion Echo Planar Imaging | Multi-Gradient Echo |
| Resolution | 0.2x0.2x0.2 mm | 0.6x0.6x0.6 mm | 0.6x0.6x0.6 mm | 0.6x0.6x0.6 mm | 0.6x0.6x0.6 mm |
| Matrix Size | 300x400x250 | 100x133x83 | 96x133x83 | 100x133x83 | 150x200x83 |
| Echo Time(s) | 6.5 ms | 6.2-198 ms (x32) | 6.2 ms | 46.5 ms | 4.6-28.2 ms (x4) |
| Repetition Time(s) | 225 ms | 1000 ms | 1612 ms | 800 ms | 70 ms |
| Flip Angle (degrees) | 30 | 90 | 90 | 90 | 15 |
| Averages | 4 | 2 | 1 | 1 | 1 |
| Total Scan Time | 18.75 hours | 4.6 hours | 4.1 hours | 20 hours | 1.8 hours |

A high-resolution anatomical (HRA) image was obtained using a Fast Low Angle Shot (FLASH) pulse sequence.

Diffusion-weighted images were acquired using a 3D Echo-Planar Imaging (EPI) pulse sequence with 6 segments. A multi-shell diffusion encoding scheme was used with b(#dirs)=0(3), 500(6), 1000(6), 1500(30), 3000(30) and 4500 (45). A second repetition of the full multi-shell acquisition was collected with the opposite phase encoding to enable geometric distortion correction in the processing stage^42^.

T2 and T1 relaxometry was conducted using Resource for Experimental Magnetic Resonance Microstructure Imaging (REMMI) pulse sequence protocols https://remmi-toolbox.github.io/, which employ the RARE pulse sequence for multiple spin echo (MSE) to support T2 and MWF mapping and MT-based selective inversion recovery (SIR) RARE to support T1 and BPF mapping ^43^.

T2* relaxometry was conducted by acquiring susceptibility-weighted images using a multi-echo 3D gradient echo (GRE) sequence.

### 2.3 Processing of Multi-parametric qMRI Data

All MRI data were transferred to the University of Arizona’s High-Performance Computing cluster and processed using various software pipelines (details and software versions in Table S1).

### 2.4 High Resolution Anatomical Template Construction

HRA images of all of the middle-age brains were used to create a HRA template using ANTs^44^ (Table S1). The HRA template served as an initial rigid for subsequent template making and for visualization of maps.

### 2.5 Diffusion MRI Processing, Mapping, Registration and Template Construction

Diffusion MRI data were processed using TORTOISE software^45,46^ (Tabl S1). Diffusion weighted MRI data were imported and reoriented to standard anatomical reference axes. Motion and distortion correction were performed including “blip-up-blip-down” correction and combination of forward and reverse phase encoded scans^47^. The diffusion tensor was fit using a nonlinear least squares algorithm. A study-specific middle-age diffusion tensor template was constructed using the DTIREG tool from TORTOISE^42,45,46^ and the remaining late-age diffusion tensors were registered to the template and DTI maps were calculated in the template space for each brain including fractional anisotropy (FA), Trace (TR) for the diffusion tensor, axial diffusivity (AD) and radial diffusivity (RD) so that all DTI maps were co-registered in this common template space for group-level analyses and SVM classifier processing.

### 2.6 Relaxometry MRI mapping and registration

MRI data were imported from bruker to nifti format using the brkraw tool ^48^ (Table S1). Metric mapping of T2 and MWF from multi-echo T2 data and mapping of BPF from multiple IR data was accomplished using the REMMI matlab toolbox https://remmi-toolbox.github.io/ (Table S1). R2* mapping from multi-echo susceptibility weighted images was performed using custom python code (Table S1). All processed maps were registered into the template space using the same registrations obtained from DTI maps.

### 2.7 ROI Segmentation for Subsequent Analyses

To assess whole brain and hippocampal volume differences between the middle and late age groups, we generated brain masks using each individual HRA brain volume. For whole brain masks, segmentation was performed semi-automatically using the active region growing contour tool in ITKSNAP^49^ (Table S1). For hippocampal masks, Regions of interest (ROIs) were manually drawn using anatomical boundaries. Volume values were then calculated from the segmentation masks and compared between the middle-age and late-age groups using a Wilcoxon test.

For all other segmentations, a rhesus macaque atlas was co-registered to the middle-age template, allowing for the extraction of morphological density percentages within each region^50^. The accuracy of the atlas registration was evaluated by comparing a manually drawn hippocampus region to the hippocampus region derived from the atlas-based region of interest (ROI) (see Figure S1).

### 2.8 Morphological Analysis

Diffusion tensor tensor-based morphometry (DT-TBM) was used to assess local volume differences between age groups^9,51^ (Table S1). To accomplish this we computed the logarithm of the determinant of the Jacobian (LogJ)^12,52,53^ for each brain in the study from the transformations generated from the diffusion tensor template registration and averaged the LogJ maps of all brains in each of the mid and late age groups (Table S1). Difference maps between late- and middle-aged brains were calculated so that positive values indicate increased local volume of the late age group compared to middle age and negative values indicate decreased local volume. This difference map was visualized by color coded overlay on the HRA middle-age template. Average LogJ values were extracted for each labeled region, and the anatomical regions were ranked by the highest to lowest (Table S1).

### 2.9 Voxel-wise Support Vector Machine Analysis of qMRI metrics

All SVMs were performed in MATLAB and subsequent analyses were performed in python using custom lab scripts (Table S1).

#### 2.9.1 All-metric, Relaxometry-only, and Diffusion-only SVM

All DTI maps – FA, TR, AD and RD – and all relaxometry maps – T2, MWF, BPF, R2* - for all brain volumes in a common template space were used to perform SVM classification of age group (Figure 2). All MRI maps were first normalized using z-scores ^54^. A support vector machine (SVM) classifier was implemented in MATLAB to predict classification accuracy at each voxel within the common template space (Table S1). The SVM was restricted to voxels within the template mask and evaluated using a 5×5×5 voxel neighborhood with 5-fold cross-validation (Table S1). To compare predictive power across qMRI techniques, three SVM models were created using training datasets from : 1) all qMRI metrics, 2) diffusion-only metrics, and 3) relaxometry-only metrics.

**Figure 2:**
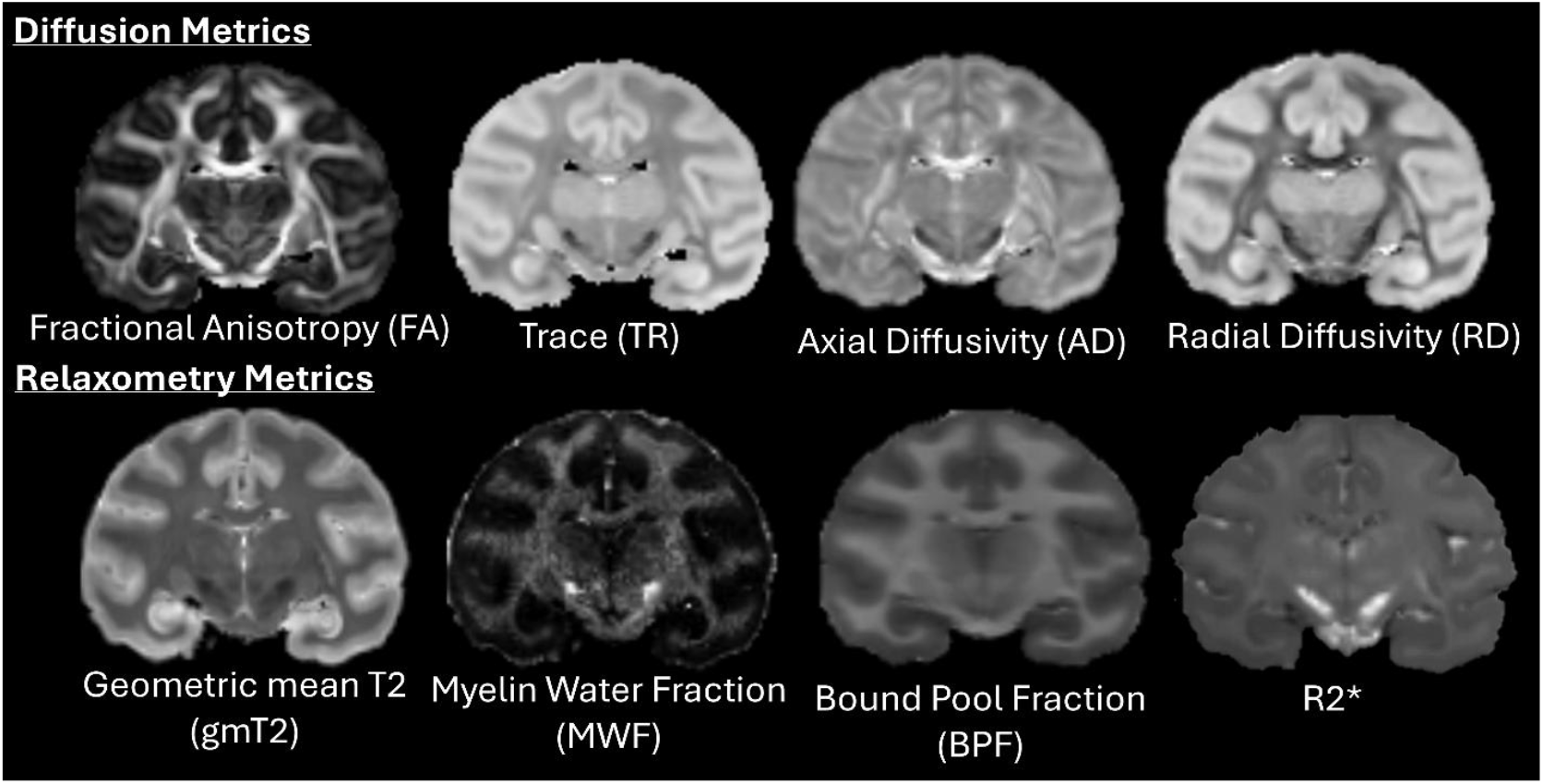
Representative Diffusion and Relaxometry Metrics - from Left to Right - FA, TR, AD,RD, T2, MWF,BPF, and R2*

Classification performance across these three models was evaluated by comparison of voxelwise accuracy values for each using density histograms and composite maps. SVM accuracy map distributions were thresholded at 95th percentile or above to capture highest performance.

A 1D histogram of the all-metric SVM accuracy values was used to determine classification regions (Table S1). Voxels exceeding 95^th^ percentile were considered, and a composite map was created from diffusion-only and non-diffusion-only SVM accuracy maps overlapping and visualized on a 3D anatomical template, with yellow=overlapping, red=non-diffusion only, and green=diffusion-only.

#### 2.9.2 Leave-One-Subject-Out Comparison to All-metric SVM

To evaluate whether a single subject disproportionately influenced classification accuracy, the all-metric SVM was repeated using a leave-one-subject-out (LOSO)^55-57^ approach, where each run omitted one subject (7 runs in total). Accuracy values were averaged across runs for each voxel and a 95th percentile threshold was applied. A ranked table of LOSO SVM was constructed for each region of interest, which showed the same relative ranking seen in all metric SVM (see Table S2).

#### 2.9.3 SVM Permutations to Evaluate Single MRI Map Contributions

In order to evaluate feature importance, “leave-one-feature-out” was performed^58,59^. To assess the contribution of individual MRI maps to classification accuracy, we performed 255 permutations of SVM training, using different combinations of MRI maps. Only permutations exceeding the 95^th^ percentile distribution were retained, and average accuracy values were computed. The impact of dataset complexity on classification, was measured via accuracy and plotted as a function of the number of MRI maps used in training (1 to 8 maps). Violin plots (Figure 8C) illustrated the contribution of individual MRI maps by comparing SVM permutations that included a specific map versus those that excluded it. The difference in 95^th^ percentile between these two conditions was computed to quantify each map’s influence on classification performance. In the region with the highest density of thresholded SVM accuracy values, the coefficients of each MRI map in the all-metric SVM were plotted to evaluate spatial coverage across different MRI map types.

## 3 Results

### 3.1 Brain Morphometric Differences Between Middle and Late Aged Brains

Age-dependent morphometry changes were mapped across the whole brain by voxelwise subtraction of average LogJ values for the late aged from the middle aged groups where positive values indicate morphometric expansion and negative values shrinkage (Figure 3). The resulting map indicated that the majority of brain regions were smaller in the late-age group compared to the middle-age group, as represented green overlay voxels (Figure 3). The cortex demonstrated prominent shrinkage across all regions although whole brain volume was not significantly different between middle and late age groups (U=11.0,p=0.143, Figure 4AB).

**Figure 3:**
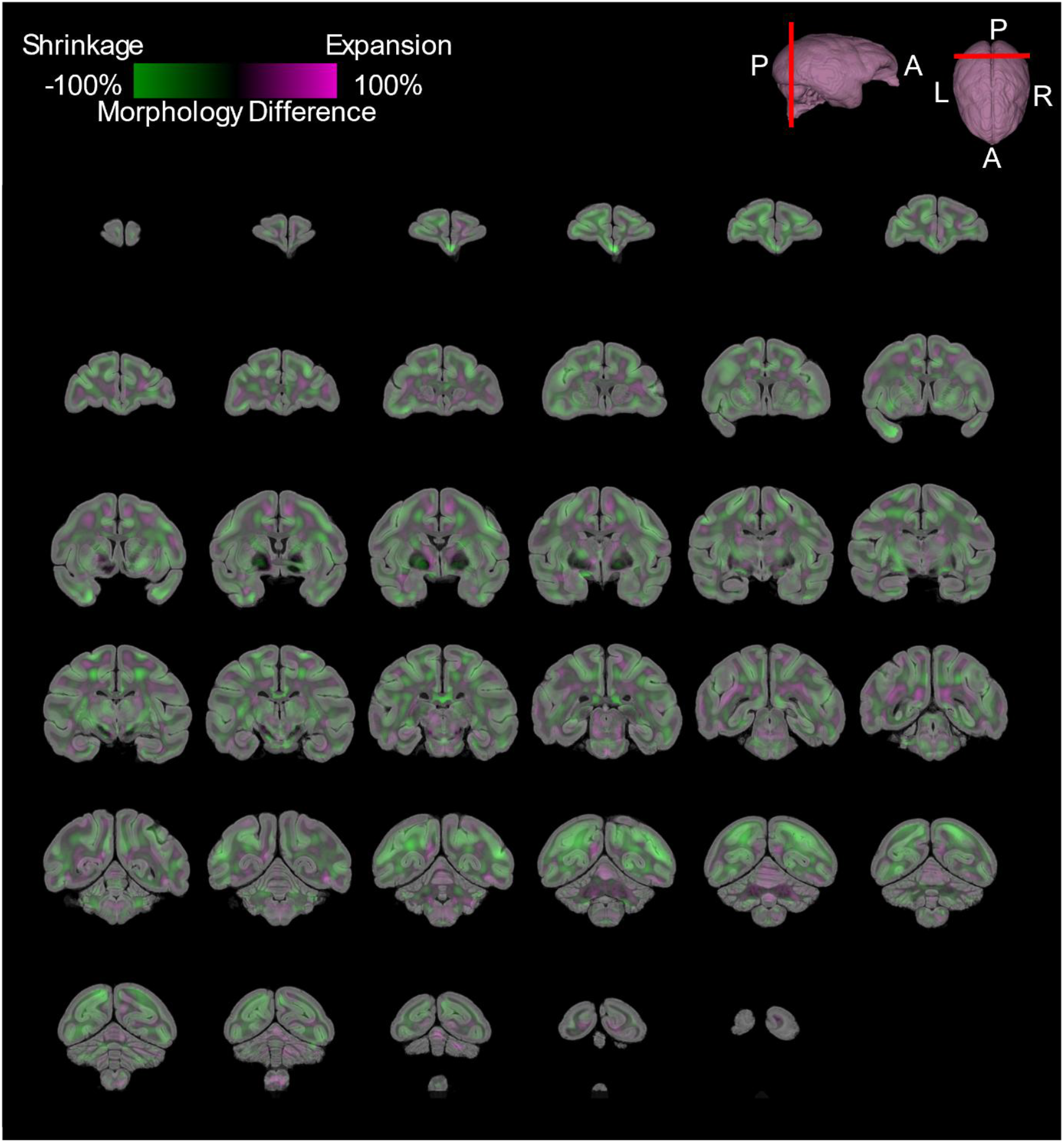
Coronal slices of Morphometry Average Difference Map Between Late Age and Middle Age. Purple colors represented -100% shrinkage and green colors represented 100% growth

**Figure 4:**
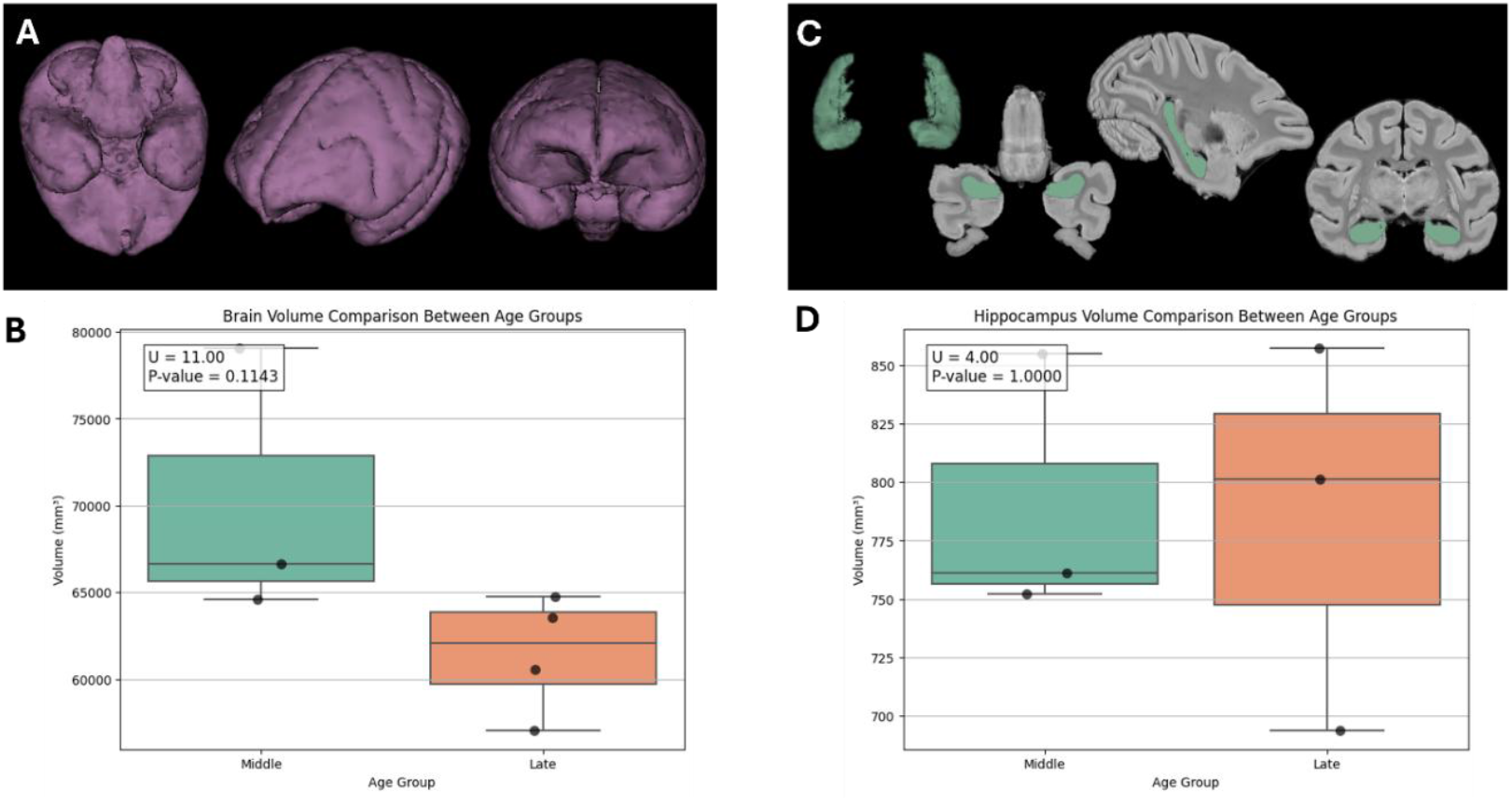
A) Representative Whole Brain Volume Renders Various Views B) Whole Brain Comparisons (Non-parametric T-test) C) Representative Hippocampal Volume Render and Segmentation Various Views D) Hippocampal Volume Comparisons (Non-parametric T-test)

Region of interest analysis of LogJ differences using atlas-defined segmentation (Table 2) found Prominent shrinkage in several neocortical regions, including the insular (−16.35%), occipital (−12.0%), frontal (−11.04%), parietal (−10.37%), cingulate (−9.77%), and orbitofrontal (−8.44%) cortices. Central white matter also showed a smaller volume in the late age group compared to the middle age by -8.72%. In contrast, regions with minimal morphological changes (<5%) included the amygdala (−3.11%) and hippocampus (−2.67%). Hippocampal volumes between middle and late age did not show a significant difference with a Wilcoxon test (U=4.0,p=1.0, Figure 4CD).There were no prominent regions of age-dependent morphometric expansion based on the brain regions selected from the atlas.

**Table 2:** Brain Regions Ranked by Largest Morphometry Differences.

| **Table 2: Brain Regions Ranked by Largest Morphometry Differences** |  |
| --- | --- |
| **Brain Region** | **Mean Logarithm of the Determinant of the Jacobian (%)** |
| Insular neocortex | -16.35+/-13.50 |
| Occipital neocortex | -12.00+/-19.45 |
| Frontal neocortex | -11.04+/-16.85 |
| Parietal neocortex | -10.37+/-15.11 |
| Striatum | -10.01+/-14.30 |
| Cingulate neocortex | -9.77+/-16.71 |
| Central white matter | -8.72+/-20.87 |
| Orbitofrontal neocortex | -8.44+/-16.56 |
| Temporal neocortex | -7.84+/-14.34 |
| Thalamus | -7.36+/-17.91 |
| Pons | -5.19+/-16.91 |
| Amygdala | -3.11+/-10.00 |
| Hippocampus | -2.67+/-12.12 |
| Septum | -1.30+/-5.32 |
| Midbrain | -1.17+/-15.10 |
| Hypothalamus | -0.085+/-12.67 |
| Cerebellum | -0.027+/-11.95 |
| Medulla | 0.034+/-12.73 |

### 3.2 Binary Classification of both Relaxometry and Diffusion MRI Metrics Yields Highest Accuracy

The classifier performance index of percent accuracy was mapped across the whole brain for three different models: Diffusion-only, Relaxometry-only and All-metric (Figure 5AB). Histogram density plots of percent accuracy values across the whole brain showed that both relaxometry-only and diffusion-only SVMs performed similarly, with the diffusion-only SVM exhibiting a longer right tail in the distribution (Figure 5B). The all-metric SVM displayed a skewed Gaussian distribution with greater median accuracy and a pronounced right-sided tail, indicating a greater number of voxels with high classification accuracies.

**Figure 5:**
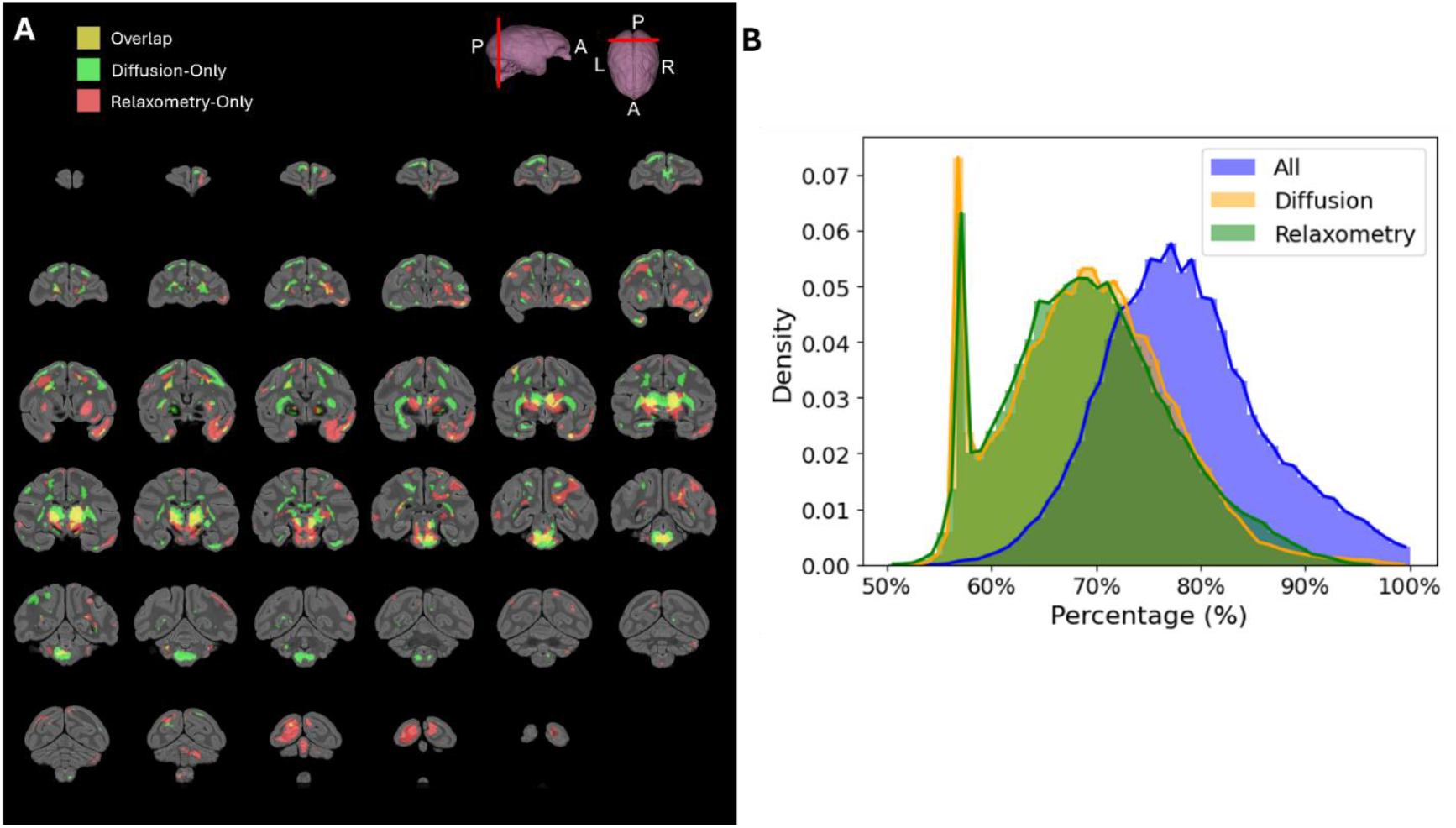
A) Composite map of overlap and non-overlap regions from diffusion and relaxometry. (>95th percentile), B) Density plot of all, diffusion-only, relaxometry-only SVMs accuracy map values

The composite map comparing the spatial distribution of diffusion-only and relaxometry-only SVM accuracy revealed distinct anatomic regions with high (>95^th^ percentile) classification accuracy (Figure 5A). There was minimal overlap (yellow regions) localized to parts of the midbrain and thalamus.

### 3.3 Highest Accuracy Regions Explained by MRI Metric Weightings

The all-metric SVM accuracy maps (Figure 6) revealed spatially distinct patterns of age classification, with the thalamus exhibiting the highest density of very good SVM performance (>95^th^ percentile) voxels (see Table 3). Coefficient maps for the all-metric SVM were used to distinguish the relative contributions of each metric to age classification and spatial distributions of individual MRI maps within the thalamus (Figure 7). The spatial distribution of very good SVM performance percentages indicated that certain regions, more than others, are more associated with age-related differences. Additionally, the coefficients of each map revealed that the SVM accuracies are spatially distinct, further indicating the individual contributions of the MRI maps on age classification performance.

**Table 3:** Brain Regions Ranked by Highest All-metrics SVM Accuracy.

| **Brain Region** | **Percentage of High SVM Accuracy Density (%)** |
| --- | --- |
| Thalamus | 61.1+/-0.46 |
| Pons | 50.25+/-0.47 |
| Midbrain | 36.96+/-0.45 |
| Striatum | 18.84+/-0.36 |
| Medulla | 16.29+/-0.34 |
| Frontal Neocortex | 9.91+/-0.27 |
| Hypothalamus | 8.88+/-0.26 |
| Central White Matter | 6.84+/-0.23 |
| Temporal Neocortex | 5.02+/-0.20 |
| Amygdala | 4.37+/-0.18 |
| Orbitofrontal Neocortex | 3.90+/-0.18 |
| Hippocampus | 2.87+/-0.15 |
| Parietal Neocortex | 2.30+/-0.14 |
| Cerebellum | 1.57+/-0.11 |
| Occipital Neocortex | 1.46+/-0.11 |
| Septum | 1.06+/-0.09 |
| Insular Neocortex | 0.89+/-0.09 |
| Cingulate Neocortex | 0.70+/-0.08 |

**Figure 6:**
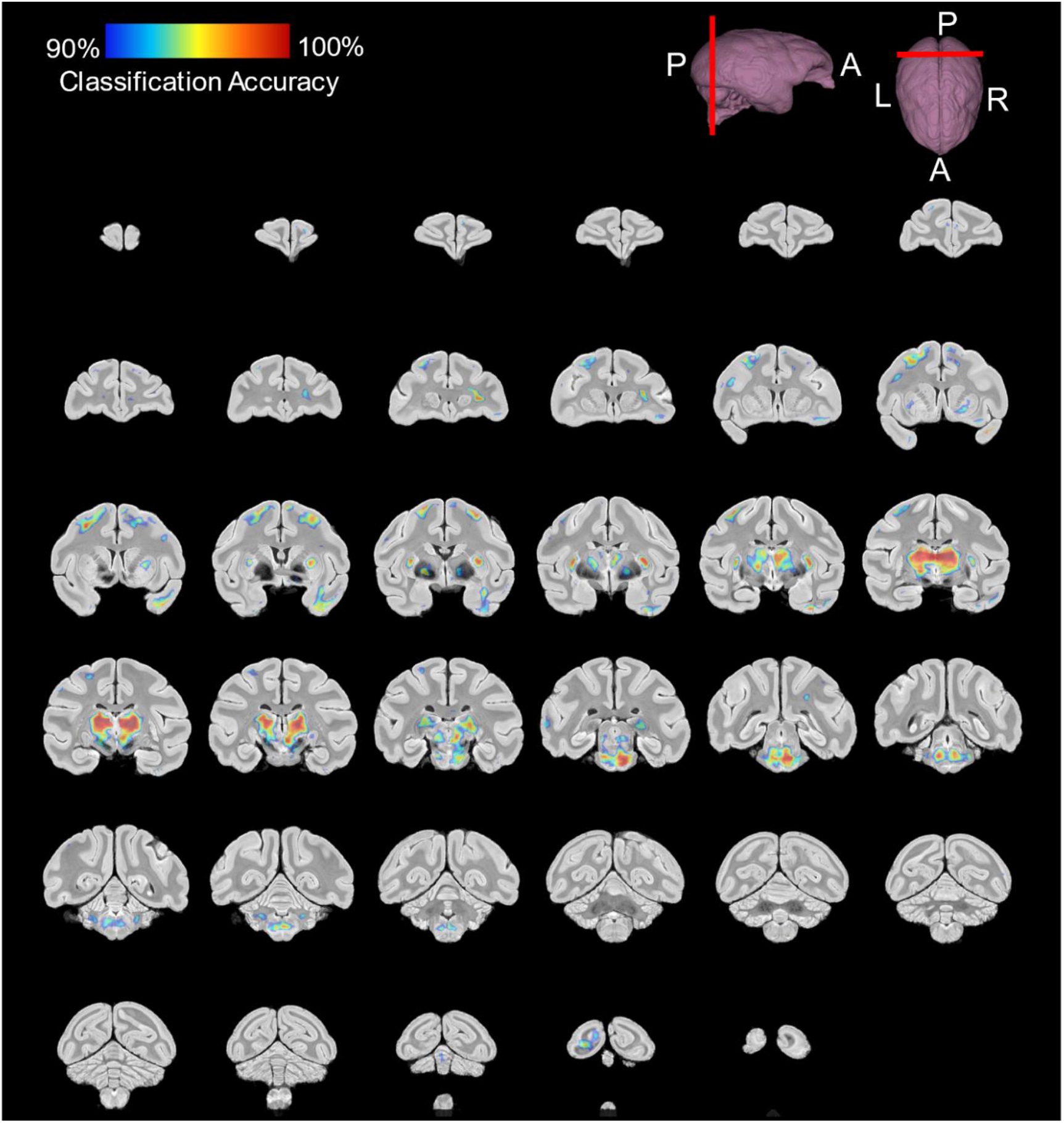
Coronal slices All metric Binary Classification Very Good Performance Accuracy Map (threshold for 95% confidence interval)

**Figure 7:**
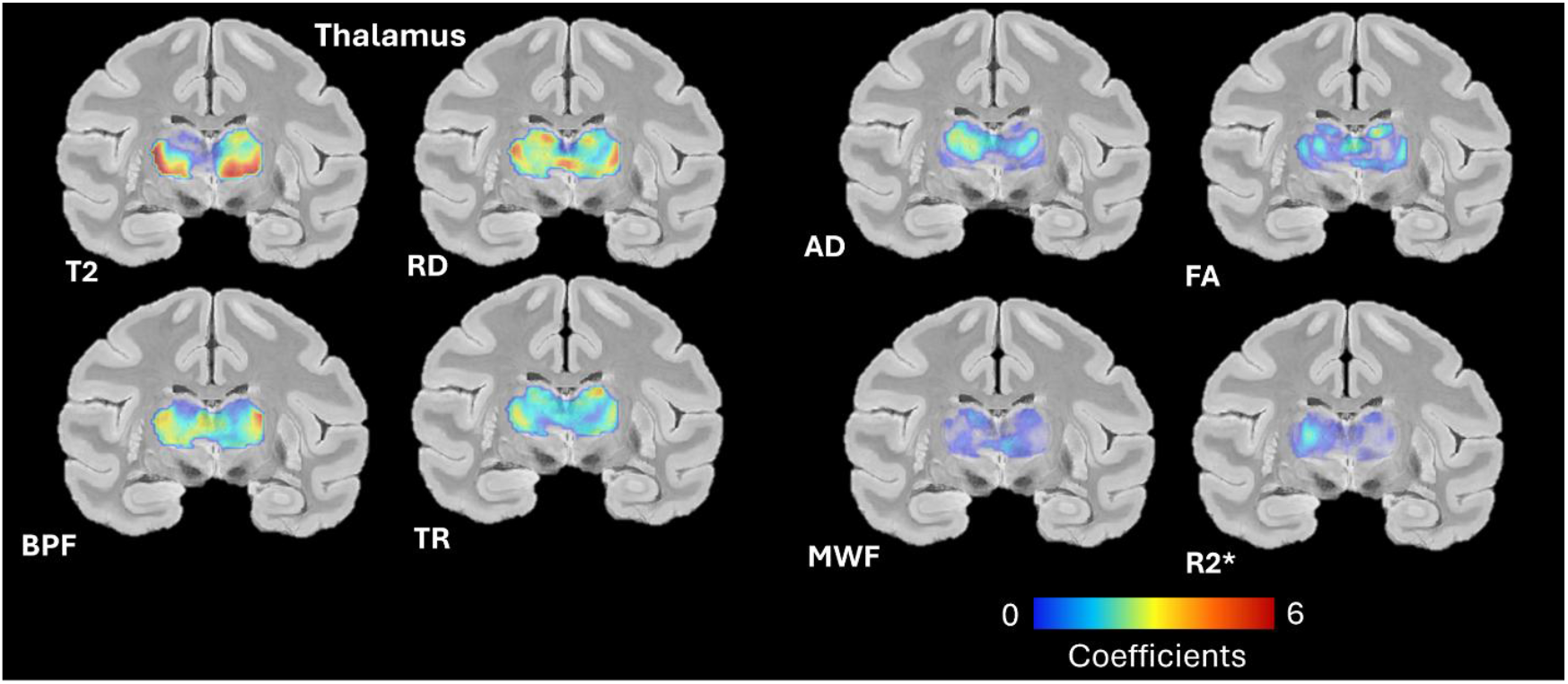
All-metric SVM Coefficients in the Thalamus

**Figure 8:**
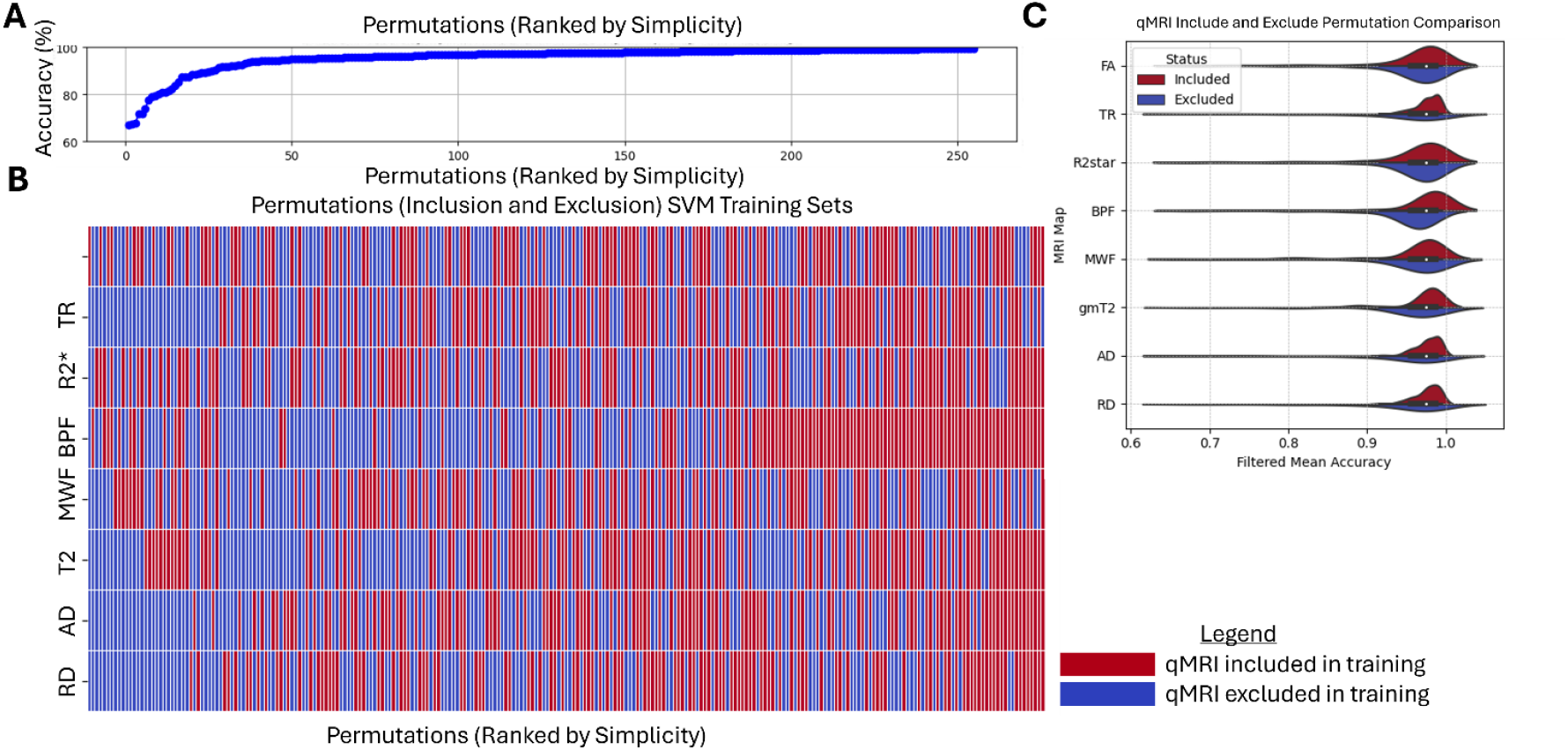
A) Permutations and their corresponding 95th percentile or above SVM Accuracy in the Thalamus B) MRI Map Permutations Ranked By Simplicity (Single Metric vs. Multiple Metrics Used for Training) C) SVM Accuracies for Each Permutation of Each Metric that Included MRI Map for Training or Excluded for Training

We ran all possible permutations that included a specific MRI map and those that excluded it and thresholded each permutation by its 95^th^ percentile or above (Figure 8AB). Inclusion-Exclusion percent difference maps were calculated for each individual MRI map (Figure 8C), we observed the following ranking for the thalamus: AD (4.01%), TR (3.99%), RD (3.79%), T2 (2.68%), BPF (2.03%), MWF (0.98%), FA (0.74%), and R2* (0.45%) (Table 4). Individually, these MRI metrics accounted for a total percent difference of 18.65% in the thalamus. Among the ranked maps, AD, TR, RD, and T2 showed a significant correlation with age.

## 4 Discussion

A data-driven, voxelwise analysis of high quality and high-resolution ex vivo qMRI maps was performed to evaluate age-dependence of morphometry, microstructure and tissue composition by region. The binary classifier for mid- or late-age classification demonstrated the highest accuracy when utilizing all MRI metrics, while classifiers using only diffusion or relaxometry data performed worse than the all-metric SVM (Figure 5B & 8A). The overall performance for diffusion-only and relaxometry-only classification were similar in distribution of percent accuracy values, but with distinct spatial patterns and the majority of predictive voxels non-overlapping between the two types of classification (Figure 5AB). Additionally, regions with large morphological differences were distinct from those with the highest age classification accuracies, suggesting that morphometry, diffusion and relaxometry metrics capture distinct tissue composition and microstructure information (Table 2 & 3). This conclusion is further supported by distinct spatial patterns of coefficient weighting for each metric in the thalamus, the region with the highest classification accuracy (Figure 7). Taken together, the findings of this study provide evidence that multi-parametric MRI can capture distinct biological features related to age-dependent brain tissue differences.

### 4.1 Preferential Morphometric Reduction in Neocortex with Age

Tensor based morphometry analysis of brains in this study revealed that the neocortex was smaller in the late-age group compared to the mid-age female bonnet macaque brains, with a 16%-7% smaller volume across the insular, occipital, frontal, cingulate, and parietal neocortex (Figure 3 and Table 2). Age-dependent neocortical shrinkage has been commonly reported in both human and animal studies {Fletcher, 2018 #301;Alexander, 2008 #43;Alexander, 2008 #489;Shamy, 2006 #454;Shamy, 2011 #455}. In the present study neocortical shrinkage was found in the absence of significant whole-brain volume differences between late and middle age, indicating selective reduction of the neocortex in late age (Figure 3 & 4B). An 8% smaller volume in central white matter was also observed between middle and late age brains. A decrease in white matter volume from middle age onward in rhesus macaques has been previously reported^61,62^. The hippocampus, a region often associated with age related atrophy, showed no significant volume differences between late- and mid-age groups (Figure 3 & 4D). Previous studies have suggested that hippocampal volume remains stable in normal aging rhesus macaques^62-66^.. One reason for this discrepancy could be the absence of comorbidities, such as hypertension or cerebrovascular disease{Raz, 2007 #327}, in controlled animal studies, which may contribute to the accelerated atrophy seen in age-susceptible regions in human studies. This highlights the importance of combining microstructural and volumetric analysis to capture more nuanced tissue compositional changes that may occur either alongside or independently of volumetric changes.

### 4.2 Thalamus: Highest Density of SVM Age Classification Accuracy

Regions found by voxelwise SVM classification to have the greatest qMRI metric alterations with age were distinct from those with morphometric change (Table 2 & 3). While regions of the neocortex and the central white matter demonstrated only modest values for accuracy, the deeper brain structures, including brain stem regions, the hypothalamus and thalamus exhibited qMRI metric alterations with age in the absence of prominent morphometric reduction (Figure 3 & 6).

Of the anatomical regions evaluated, the thalamus contained the most voxels with very good SVM performance, making it the most age-sensitive region distinguishing late and mid-age female bonnet macaque brains (Figure 6). No single qMRI metric was responsible for a large percentage of SVM accuracy performance – with more metrics included in training ultimately improving accuracy in the thalamus (Figure 8A). This finding suggests that individual qMRI metrics likely capture complementary features of tissue composition and microstructure which may be different at late and mid-age. The ranking of qMRI metrics within the thalamus based on their relative weights in SVM accuracy showed that diffusivity metrics (AD, RD and TR) were most consequential followed by T2 values (Table 4 and Figure S2). These four metrics were also found to have significant correlation with age while the remaining metrics (BPF, MWF, FA, and R2*) appeared to be less consequential for age classification and did not demonstrate correlation with age. Previous literature has alsosuggested that RD and TR increase in the thalamus with age, however it is unknown as to the exact cause for these metrics to increase, it suggested that it may be due to decrease in the number of myelinated fibers present in the white matter regions of the thalamus^67^. Additionally, in vivo human MRI studies have shown significant positive correlations between T2 values and age^68,69^. Taken together, these findings imply that differences in diffusion metrics (AD, TR, RD) reflect reduced structural barriers, resulting in increased diffusion in all directions and higher metric values. The increase in T2 may be similarly indicative of increased water content due to reduction of cell barriers. Another interpretation is decreased iron content, although the lack of findings related to R2*, (Figure 8C & S2 & Table 5) values in this study suggest that this is unlikely.

**Table 5:** Feature Importance Analysis Using Leave-One-Fature-Out

| Order of Difference between Inclusion and Exclusion | Percent Difference Between Inclusion and Exclusion SVM Accuracy |  | R Value (* p<0.05) |
| --- | --- | --- | --- |
| 1 | 4.01% | AD | **0.86\*** |
| 2 | 3.99% | TR | **0.86\*** |
| 3 | 3.79% | RD | **0.86\*** |
| 4 | 2.68% | T2 | **0.82\*** |
| 5 | 2.03% | BPF | 0.46 |
| 6 | 0.98% | MWF | 0.75 |
| 7 | 0.73% | FA | 0.32 |
| 8 | 0.45% | R2* | 0.5 |

### 4.3 Diffusion and Relaxometry Metrics Provide Distinct Information about Age-Related Differences

Our study suggests a combinatorial effect when assessing quantitative MRI maps, which may not be evident in univariate approaches. Multi-variate approaches, such as SVM classifiers, are advantageous for probing multiple biological processes associated with aging, such as iron deposition and changes in the composition and microstructure of grey and white matter.

Expanding the SVM approach to a voxel-wise analysis in this study offers a bias-free assessment of spatially dependent tissue differences in the brain, and identified regions such as the thalamus with the most prominent age-dependent changes in the combined metrics (Figure 6). In addition to identifying age-prone regions, voxel-wise SVMs can aid in designing qMRI protocols and batteries that are informative of aging without redundancy.

The findings of the current study suggest that evaluation that relies solely on one class of quantitative MRI maps, such as diffusion or relaxometry, may have limited sensitivity and specificity regarding tissue structure. Our diffusion-only and relaxometry-only SVMs highlight this limitation, as they covered distinct spatial regions with minimal overlap (Figure 5AB). In addition, each was found to have an overall lower SVM accuracy compared to that of the all-metric SVM. This evidence reinforces the idea that using MRI metrics from both categories provides the most comprehensive information regarding age-related differences.

## 5 Limitations and Future Directions

This study was limited by a small sample size, as it focused on a precious, but small cohort of bonnet macaques; and this may have contributed to variability in permutation test accuracy for certain quantitative MRI maps. However, the leave-one-out subject analysis confirmed that classification accuracy was not driven by a single subject but reflected genuine differences between the middle- and late-age brains, increasing confidence in the SVM approach for this cohort. The same highest ranking of regions of the highest density of SVM accuracies in the leave-one subject out and all metric SVM further confirmed this approach (see Table S2).

Additionally, SVMs are known to have balanced predictive performance even in applications with small datasets compared to other approaches^4^. As open-access repositories and data-sharing initiatives expand, our binary classification approach, well-suited for large datasets, can be applied to broader populations. For longitudinal studies, transitioning from a binary to a multi-classifier model may enhance applicability. While our SVM approach identifies brain regions susceptible to aging, future research should explore approaches that assess the directionality of these changes. Our SVM results provide an initial evaluation of this cohort, which can be validated through ground-truth methods such as immunohistochemistry (IHC) to confirm observed structural differences. Finally, because *ex vivo* imaging requires tissue preparation, our findings should be interpreted with caution when generalizing to *in vivo* research.

## Data and Code Availability

All code used for analysis is available here: https://osf.io/w9ck3/?view_only=f3611ea0fc854de395f78216ee4da7b5. Code and protocols will be made publicly available on github: https://github.com/UAmsbil upon publication. All raw data and github repository code used in this study will be made available via the University of Arizona upon publication.

## Declaration of Competing Interest

There are no conflicts of interests for any author to declare.

## Author Contributions

Laurel Dieckhaus: Data curation, diffusion and relaxometry MRI processing, analysis and methodology, data interpretation and writing (drafts, editing, figures). Kelsey McDermott: data interpretation and writing (reviewing and editing). Jean-Philippe Galons: Expertise in MRI acquisition, aided in troubleshooting data curation process and data interpretation. Carol Barnes: Expertise in bonnet macaque aging, conceptualization, specimen curation, and writing (reviewing and editing). Elizabeth B. Hutchinson: Conceptualization, funding acquisition, methodology and resources, project supervision, interpretation and writing (reviewing and editing).

## Acknowledgements

This work was generously funded and supported by the McKnight Brain Research Foundation, Arizona Alzheimer’s Consortium, Department of Health, UA translational bioimaging resource (TBIR), NIH small instrumentation grant (**S10 OD025016**), State of Arizona. Research was also supported by the National Institute of Aging of the National Institutes of Health under award number **T32AG082631-01**. High Performance Computing (HPC) resources supported by the University of Arizona TRIF, UITS, and Research, Innovation, and Impact (RII) and maintained by the UArizona Research Technologies department. We are grateful to the UA KEYS program for student training and funding (ZA), in which Avinash Murlikrishnan, a KEYs student who aided in the creation of the hippocampus ROIs used in the manuscript, would have not been possible. The authors would like to thank Michael Cardenas for feedback on the Support Vector Machine implementation. Additionally, the authors would like to thank Leili Falbinan, who helped to organize the github repositories and code protocols referenced in this manuscript. Also, the generous time that was dedicated from Jean-Philippe Galons at the MRI scanner to troubleshoot acquisition issues and helped to make the Diffusion Tensor Imaging acquisition possible. Also, Kevin Harkins, who made the REMMI acquisition feasible on our MR scanner.

## Funding Sources

This work was generously funded and supported by the: McKnight Brain Research Foundation, Arizona Alzheimer’s Consortium, UA translational bioimaging resource (TBIR) and NIH small instrumentation grant (**S10 OD025016**), State of Arizona. Research was also supported by the National Institute of Aging of the National Institutes of Health under award number **T32AG082631-01**.

## Notes

### Competing Interest Statement

The authors have declared no competing interest.

