## Supplemental Figures and Tables for "Multi-Metric Quantitative MRI Identifies Spatially Distinct Age-Related Brain Differences in Female Bonnet Macaques"

| Map Type/Procedure | Source Code or Reference | Repository |
|---|---|---|
| MRI-Compatible Container | Lab-based protocol | https://github.com/UAmsbil/Lab3DModels/tree/main/MRI/Tissue_Holders |
| Ex vivo tissue preparation | Lab-based protocol | https://github.com/UAmsbil/WetLabPrep/tree/main/ExVivo_Prep |
| Bruker Raw Data Importation | Brkraw | https://github.com/BrkRaw |
| High Resolution Anatomical | ANTs | https://github.com/ANTsX/ANTs |
| T2 | REMMI | https://github.com/remmi-toolbox/remmi-matlab |
| MWF | REMMI | https://github.com/remmi-toolbox/remmi-matlab |
| T1 | REMMI | https://github.com/remmi-toolbox/remmi-matlab |
| BPF | REMMI | https://github.com/remmi-toolbox/remmi-matlab |
| DTI | TORTOISE V3.2 | https://tortoise.nibib.nih.gov/ |
| R2* | Lab-written code | https://github.com/UAmsbil/Quantitative_MRI/tree/main?tab=readme-ov-file |
| Region of Interest and Segmentation | ITKSNAP | |
| Initial Rigid Template Creation | ANTs | https://github.com/ANTsX/ANTs |
| Diffusion-Tensor Tensor Based Morphometry | TORTOISE V3.2 | https://tortoise.nibib.nih.gov/ |
| Volume Analysis | Lab-written code | https://github.com/UAmsbil/MRIAnalysis/tree/main/VolumeAnalysis |
| Morphology Subtraction Map | Lab-written code | https://github.com/UAmsbil/MRIAnalysis/tree/main/SubtractionMaps |
| Morphology Table Ranking | Lab-written code | https://github.com/UAmsbil/MRIAnalysis/tree/main/Morphology_LogJ |
| Support Vector Machine | Lab-written code | https://github.com/UAmsbil/MRIAnalysis/tree/main/SVM |
| SVM Permutation Figures in Thalamus | Lab-written code | https://github.com/UAmsbil/MRIAnalysis/tree/main/VisualAids |
| Histograms | Lab-written code | https://github.com/UAmsbil/MRIAnalysis/tree/main/VisualAids |

**Table S1: Code Repositories for Acquisition, Processing Raw MRI Data, and Analysis**

*Table S1: Details of Software and its corresponding versions for Image Processing and Analysis*

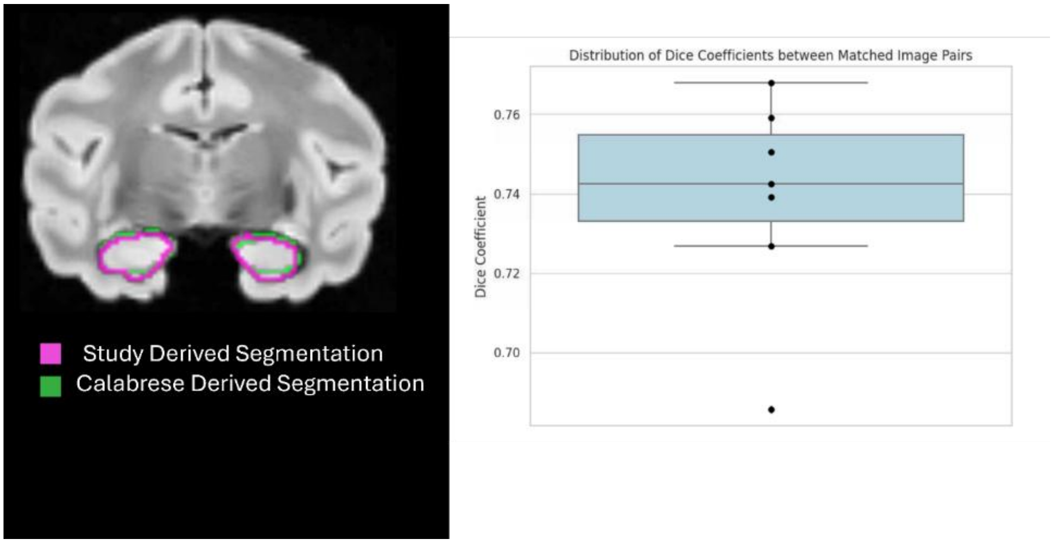


*Figure S1: Hippocampus DICE coefficients compared to atlas A) Representative overlay of atlas on study specific segmentation*

**Table S2: Brain Regions Ranked by Highest Leave-One-Subject-Out All-metrics SVM Accuracy**

| Brain Region | Percentage of High SVM Accuracy Density (%) |
|---|---|
| Thalamus | 58.32+/-0.47 |
| Pons | 50.27+/-0.47 |
| Midbrain | 34.24+/-0.44 |
| Striatum | 19.20+/-0.37 |
| Medulla | 16.79+/-0.35 |
| Frontal Neocortex | 9.75+/-0.28 |
| Amygdala | 8.32+/-0.25 |
| Hypothalamus | 7.30+/-0.24 |
| Central White Matter | 6.44+/-0.23 |
| Temporal Neocortex | 5.04+/-0.20 |
| Orbitofrontal Neocortex | 3.68+/-0.17 |
| Hippocampus | 2.55+/-0.14 |
| Parietal Neocortex | 1.80+/-0.122 |
| Cerebellum | 1.44+/-0.11 |
| Occipital Neocortex | 1.18+/-0.10 |
| Insular Neocortex | 0.92+/-0.09 |
| Septum | 0.76+/-0.08 |
| Cingulate Neocortex | 0.66+/-0.07 |

*Table S2: Leave One Subject Out SVM Anatomical Region Ranking*

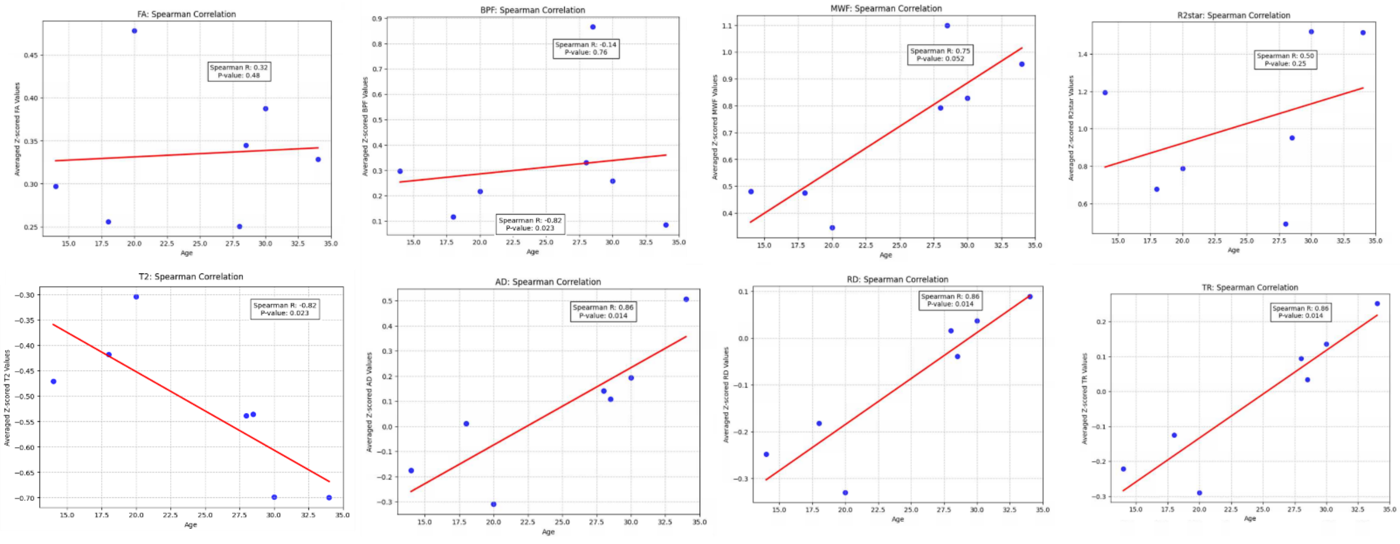


*Figure S2: Thalamus Spearman Correlations*
